# Roles of histone chaperone Nap1 and histone acetylation in regulating phase-separation of chromatin arrays

**DOI:** 10.1101/2025.05.09.653121

**Authors:** Jia Gao, Hongyun Li, Song Tan, Ruobo Zhou, Tae-Hee Lee

## Abstract

Chromatin condensation is dynamically regulated throughout the cell cycle and plays key roles in modulating gene accessibility at the highest physical level in a cell. The DNA-histone dynamics in short- and long-range inter-nucleosomal interactions are central to the regulation mechanisms of chromatin condensation which remain under active investigation. We employed 12-mer nucleosome arrays to investigate the roles of histone chaperone Nap1, and histone H3 and H4 N-terminal tails and their acetylation in regulating the formation and the properties of phase-separated chromatin condensates. These arrays form liquid-like condensate droplets under a physiological salt condition of 150 mM NaCl. According to our results from bright-field microscopy, fluorescence recovery after photobleaching, optical super-resolution imaging, and microrheology with optical tweezers, histone H4 tail lysine residues are the main drivers of liquid-liquid phase separation of chromatin arrays. We also found that the condensed liquid-like droplets contain both a mobile fraction and a relatively immobile structural scaffold and that histone chaperone Nap1 and histone H3 tail acetylation facilitate DNA-histone dynamics within the structural scaffold to lower the overall viscosity of the droplets. These results suggest that histone chaperone and histone H3/H4 tails play critical roles in regulating chromatin condensation and gene accessibility in condensed chromatin.

## Introduction

The eukaryotic cell nucleus consists of two main compartments, one of which is the chromatin that contains the genome. Although not separated by a physical barrier, chromatin exists in its own phase, while the nucleoplasm can also contain other membrane-less compartments and liquid-like droplets^1–3^. The structure of chromatin is formed largely by arrays of genomic DNA, histone proteins, and various other chromatin-associated proteins. The fundamental gene packing unit in chromatin is the nucleosome, which is ∼150 bp DNA wrapped around an octameric histone core containing two H2A-H2B dimers and one (H3-H4)_2_ tetramer^4^. The concentration of nucleosomes in a cell nucleus is in the range of a few tens to hundreds of μM^5^. The interactions between DNA and histones, both within and across nucleosomes, are central to the often complex mechanisms of gene regulation at multiple levels^6–8^. One way to regulate such dynamic genome activities is through histone post-translational modifications^9–11^. Another involves enzymes that act on chromatin, such as histone chaperones and chromatin remodelers^12–15^.

Efforts at multiple levels both *in vivo* and *in vitro* are being expended on elucidating the mechanisms of how these factors help regulate genome activity^16–22^. In particular, *in vitro* approaches with highly refined systems have made valuable contributions to deciphering complex changes at the nucleosome and DNA-histone levels without interference from unknown factors^23–26^. Many such approaches employ observing and controlling nucleosomes at the single-nucleosome level, either in individual nucleosomes or in nucleosomes linked in an array.

However, these systems lack a fundamental aspect of chromatin where nucleosomes are densely packed at a high concentration. Once the concentration of nucleosomes or nucleosome arrays is elevated to more than a few tens to hundreds of nM under a physiological ionic condition, they undergo liquid-liquid phase separation (LLPS) to form liquid-like droplet condensates^27,28^. The concentrations of nucleosomes within these droplets match those *in vivo*^27^. While the mechanism of condensed droplet formation remains under active investigation, it should involve both short- and long-range inter-nucleosomal interactions, which are likely relevant to chromatin condensation *in vivo*. A previous paper reported that the droplets are more gel-like than liquid-like, although the shape of many droplets is still quite spherical, suggesting that the droplets begin in a liquid-like state and age into a gel-like state^29^. Nevertheless, these condensates, especially those made of nucleosome arrays, are excellent platforms for investigating the physical and biological properties and functions of nucleosomes and chromatin-acting enzymes in a chromatin-like environment.

Histone N-terminal tail acetylations on H3 or H4 have been implicated in weakened intra- and inter-nucleosomal DNA-histone interactions^30–35^, thereby modulating the strength of chromatin condensation at both the nucleosome and chromatin levels. Histone H3 tail acetylation is more strongly associated with intra-nucleosomal DNA-histone compaction, while H4 tail acetylation is more strongly coupled to inter-nucleosomal chromatin compaction^30–35^. Previous reports suggested that histone acetylation generally inhibits phase separation of nucleosomes or nucleosome arrays, or that it reduces droplet size or causes them to dissolve^27,36^. However, no studies have reported how specific histone tail acetylation affects the condensation behavior of nucleosome arrays. Another important mediator of DNA-histone interactions is the class of histone chaperones. Histone chaperones contain acidic residues that can compete for histone binding against DNA^37^. Histone chaperone Nap1 can bind both H2A-H2B and (H3-H4)_2_ in a stoichiometric manner^38,39^. Nap1 mediates DNA-histone interactions to bring a random mixture of DNA and histones to their thermodynamically favored state which is the nucleosome^40^.

Despite its important role in regulating nucleosome stability, it has never been reported how Nap1 functions in a condensed phase of nucleosomes. In particular, it remains unknown on what timescale Nap1 enables or facilitates DNA-histone dynamics to eventually stabilize dynamic nucleosomes in a condensed phase. To this end, we investigated how histone H3 or H4 acetylation and histone chaperone Nap1 regulate the formation and the properties of liquid-like chromatin condensates spontaneously formed by LLPS of 12-mer nucleosome arrays. We found that H4 tail acetylation inhibits chromatin condensation while an H3 tail acetylation mimic (lysine residues mutated to glutamine) makes the condensed chromatin droplets more fluidic, based on bright-field microscopic imaging, fluorescence recovery after photobleaching (FRAP) observations, and microrheological measurements with optical traps. We also discovered that histone chaperone Nap1 lowers the viscosity of the chromatin droplets, while it significantly elevates the nucleosome concentration within a condensate. Optical super-resolution images revealed that condensed droplets contain both mobile and relatively immobile structural scaffold nucleosome arrays. The microrheology results indicate that the effects of the H3 tail acetylation mimic and Nap1 are exerted specifically on the nucleosome arrays associated with the structural scaffold. In summary, our study reports that histone H4 tail acetylation inhibits chromatin condensation, while histone H3 tail acetylation and histone chaperone Nap1 render a more fluidic yet stable structure of condensed chromatin, suggesting their roles in regulating gene accessibility at a chromatin level.

## Methods

### DNA purification

The plasmid containing 12 repeats of the Widom 601 sequence with a 25 bp linker length, pWM+12×601_172NRL, was a generous gift from the Rosen lab. The 12×601 array DNA was prepared largely as previously described^27^. Briefly, we transformed the plasmid into Top 10 Competent Cells (Thermo Fisher Scientific, Waltham, MA) and plated it on LB agar plates supplemented with 100 μg/mL carbenicillin for growth overnight. After inoculated with a small-scale preculture from a single colony, a 6 L of LB culture with 100 μg/mL carbenicillin was shaker-incubated overnight at 37 °C. The culture was harvested by centrifugation, and the plasmid was purified with the Plasmid Maxi kit (Qiagen, Hilden, Germany). The plasmid was digested with restriction enzyme EcoRV (New England Biolabs, Ipswich, MA), and then the 12×601 array DNA was purified through size exclusion chromatography (Sephacryl S-500 HR, Cytiva, Netherlands) as shown in Fig. S1. The purification step also generated competitor DNA fragments (∼300 bp) used for later nucleosome array preparation (Fig. S1).

To attach a biotin molecule at one end of the DNA, a plasmid containing 12 repeats of the Widom 601 sequence with a 25 bp linker length^41^ (a gift from Dr. Sergei Grigoryev, the Pennsylvania State University College of Medicine) was digested with BamHI, HindIII, HaeII, and DraI (New England Biolabs) to leave the nucleosome array DNA with a 4-nucleotide overhang at both ends. The HaeII and DraI enzymes were to digest the plasmid backbone into shorter fragments. The digested plasmid was purified using the same protocols described above. Two short adapter DNA fragments (Integrated DNA Technologies, Coralville, IA) were ligated to the sticky ends of the purified nucleosome array DNA with T4 DNA ligase (New England Biolabs) at 16 °C for 14 hours followed by inactivation at 65 °C for 20 minutes. The sequences of the adapter DNA fragments are phos-GATCCAGTACCTAGCATT-biotin and CTAGGTACTG for the BamHI end, and phos-AGCTTAAGCTGAGT and ACTCAGCTTA for the HindIII end, where “phos-” indicates 5’ phosphorylation. The resulting DNA has one end labeled with a biotin molecule and was purified with a PCR clean-up kit (Qiagen). The purity was confirmed with a 1 % agarose gel which showed a single band at the appropriate molecular weight.

### Nucleosome array assembly and purification

Wild-type human histones and other mutants (tailless histones, acetylation mimics, and histone octamers containing H4E63C/H3C96S C110A or H2BT115C/H3C96S C110A triple-mutations for fluorophore labeling) were purchased from the Histone Source (Colorado State University). For histone fluorophore-labeling, histones were incubated with a 10x molar excess of maleimide-functionalized Alexa Fluor 647 (Thermo Fisher Scientific) in a 10 mM Tris-HCl buffer (pH 7.5) for 2 hours at room temperature, followed by dialysis to remove unreacted fluorophores. As previously described, 12-mer nucleosome arrays were reconstituted by dialyzing a mixture of histones and nucleosomal DNA at a fixed ratio in the presence of competitor DNA fragments in a dialysis device (Slide-A-Lyzer MINI Dialysis Device, 3.5K MWCO, Thermo Fisher Scientific) against 1x TE (10 mM Tris-HCl, 1 mM EDTA, pH 8.0) buffer and stepwise decreasing salt concentrations of 2000, 1500, 1000, 750, 500, and 5 mM NaCl stepwise^42^. The ratio of DNA:histone was determined by titrating histone to DNA to ensure saturation of histone to nucleosomal DNA (Fig. S2). The assembled arrays were fractionated via size exclusion chromatography (Sephacryl S500-HR, Cytiva) (Fig. S3). The quality of each fraction was assessed by TEM imaging (for arrays with no modifications, Fig. S4) and analytical ultracentrifugation (Fig. S3). The fraction with the highest purity was selected for experiments.

### Nap1 purification

6xHis-Yeast nucleosome assembly protein 1 (Nap1) was expressed in *E. coli* and purified with Ni-NTA beads (Thermo Fisher Scientific) as reported in a previous publication^43^.

### Purification of Piccolo NuA4 and Ada2/Ada3/Gcn5 HAT complexes

Full length Esa1, Yng2(1–218), and hexahistidine-tagged Epl1(51–380) subunits of yeast Piccolo NuA4 were coexpressed using the pST44 polycistronic expression vector in BL21(DE3)pLysS cells as described previously^44,45^. The Piccolo NuA4 complex was purified by Talon (Clontech) cobalt metal affinity chromatography, followed by removal of the hexahistidine tag by TEV protease digestion and subsequent SourceQ (Cytiva) anion-exchange chromatography.

The yeast Ada2/Ada3/Gcn5 complex containing full length Ada2 residues, C-terminally hexahistidine tagged Ada3(187-702) and full length Gcn5 was coexpressed using the pST44 polycistronic expression vector^45^. Protein expression in BL21(DE3)pLysS host strains was induced with 0.2 mM IPTG at 18 °C. The Ada2/Ada3/Gcn5 complex was purified from cell lysates by Talon (Clontech) cobalt metal affinity chromatography, followed by SourceQ (Cytiva) cation-exchange chromatography.

### Phase separation procedure

Nucleosome arrays were stored in the final dialysis buffer (5 mM NaCl, 1X TE, 1 mM Dithiothreitol). The addition of the phase separation buffer resulted in the final mixture composition of 150 mM NaCl, 25 mM Tris-HCl (pH 7.5), and 200 nM nucleosome arrays. After 30 minutes of incubation, the mixture was transferred to fluidic channels constructed on a microscope slide surface, which had been passivated with methoxy poly-ethyleneglycol (mPEG, 2kDa molecular weight, Laysan Bio, Arab, AL) and bovine serum albumin (BSA, Millipore Sigma). Detailed methods of surface treatment and fluidic channel construction can be found elsewhere^34,35,46^.

### Bright field imaging

Bright field microscope images were taken on a Nikon Eclipse TE2000 inverted microscope with a Nikon Plan Apo 60x water-immersion objective. The pictures were recorded with an IAI CVM50 Industrial CCD Camera (147×114 μm^2^, 752 x 582 pixels with a pixel size of 196 nm).

The protocols for microscope slide preparation have been reported elsewhere^47^.

### Fluorescence Recovery After Photobleaching (FRAP)

Confocal fluorescence images were captured on a Zeiss LSM 880 confocal laser scanning microscope equipped with a 63X 1.4 NA objective lens and 34-channel NLO array detectors. Fluorescently labeled arrays were mixed with unlabeled arrays and biotinylated arrays at a 2(fluorescent):7(non-fluorescent):1(biotinylated) molar ratio. Biotinylated arrays were mixed to immobilize droplets on a coverslip. A surface-passivated coverslip was used for imaging. The coverslip was coated with mPEG (2k Molecular weight, Laysan Bio, Arab, AL) and biotin-PEG (5k Molecular weight, Laysan Bio, Arab, AL). Detailed surface preparation protocols can be found elsewhere^34,35,46^. Streptavidin (0.2 mg/mL, Thermo Fisher Scientific) was first incubated on a coverslip and rinsed out with the phase separation buffer before droplets were loaded.

Fluorescence Recovery After Photobleaching (FRAP) was achieved with the built-in bleaching function in the ZEN software package provided by the microscope manufacturer (Zeiss, Germany). A normalized fluorescence recovery trajectory was fit to an exponential function 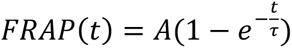 where τ is the recovery time and A is the recovery fraction.

### Microrheology measurements with optical traps

A C-trap optical tweezers instrument (Lumicks, Netherlands) was used to probe the viscoelasticity of a droplet by oscillating an optically trapped polystyrene bead (monodisperse analytical standard, 1 μm diameter, Supelco® via Millipore Sigma) inside. While the bead was oscillating, the trap position (i.e., bead position) and the force feedback were recorded. The temperature of the stage and the sample chamber were maintained at 25 °C during the measurements. The analysis followed previous publications^48,49^. Briefly, the trajectories of the bead position (X) and the force feedback (F) were fit to the following two equations to extract the elastic and viscous moduli.

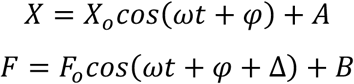

where 𝜔 is the angular frequency of bead oscillation, 𝑋*_o_* 𝑎𝑛𝑑 𝐹*_o_* are the amplitudes of the cosine waves representing the bead position and the force feedback, respectively, 𝜑 is a phase offset of the bead position oscillation, Δ is the phase lag between the bead position and the force feedback, and A and B are the amplitude offsets for the bead position and the force feedback, respectively. Once the values of 𝜔, 𝜑, 𝑋*_o_*, 𝐹*_o_*, 𝑎𝑛𝑑 Δ were determined from the fitting, the following equations were used to compute the elastic (G’) and viscous (G”) moduli at the frequency (𝜔). The force constant of the optical trap (*k*) was acquired prior to each experiment.

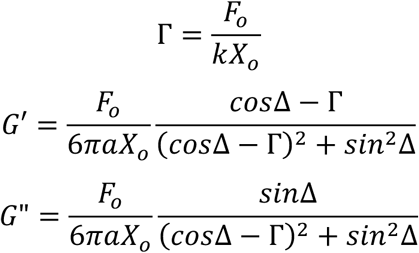

The obtained moduli values at 7 different frequencies were fit to a Burgers material model with two Maxwell components where the two relaxation components are linearly combined. The equations used for fitting are as follows. The values of τ and η were obtained from the fitting.

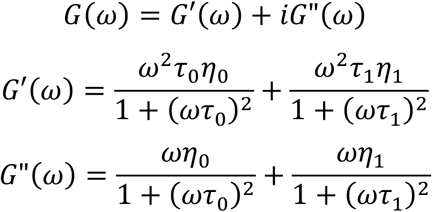

where τ and η are the relaxation time and viscosity, respectively, and the subscripts 0 and 1 are the indices of the two components. The two linearly combined relaxation components are more explicit in the time domain modulus as follows.

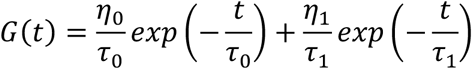

The elastic moduli for a sudden and sustained strain are the amplitudes of the two components as following.

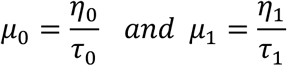

### Stochastic Optical Reconstruction Microscopy (STORM)

Three-dimensional (3D) STORM imaging was performed using a custom-built total internal reflection fluorescence (TIRF) microscopy based on a Nikon Eclipse Ti2-U inverted microscope body. A 642-nm laser (MPB Communications, 2RU-VFL-P-2000-642-B1R) was used for excitation of Alexa Fluor 647, while a 405-nm laser (Coherent, OBIS 405 nm LX) was used to reactivate fluorophores from the dark state during imaging. Both lasers were introduced through the microscopés back port and directed into a high numerical aperture (NA = 1.45) 100x oil-immersion objective via a multiband dichroic mirror (ZT405/488/561/640rpcv2, Chroma). Fluorescence emission was filtered using a quad-band emission filter (ZET405/488/561/640m-TRFv2, Chroma) and detected by an EMCCD camera (iXon Life 897, Andor Technology). A TIRF lens mounted on a translation stage allowed lateral displacement of the laser beams at the objective’s back aperture, ensuring illumination at angles just below the critical angle of the glass– water interface and selectively exciting fluorophores within ∼2 µm of the coverslip surface. A cylindrical lens (f = 1000 mm) was introduced into the emission path to introduce axial astigmatism, causing the point spread function (PSF) of single-molecule fluorescence to appear elliptical when emitters were above or below the focal plane. This PSF ellipticity was used to determine the z-position of each molecule, enabling 3D STORM reconstruction^50^. The astigmatic single-molecule images were fitted with an elliptical Gaussian function to extract the centroid in the x–y plane, and the z-position was computed based on the ellipticity of the PSF, which systematically varies with axial displacement from the focal plane.

For STORM imaging, the chromatin condensate samples were mounted in an imaging buffer containing 25 mM Tris-HCl (pH 7.5), 150 mM NaCl, 5% (w/v) glucose, 10 mM cysteamine (MEA), 0.8 mg/mL glucose oxidase, and 40 μg/mL catalase. The condensates were formed with a mixture of 1:500 fluorescently labeled:unlabeled nucleosome arrays. The fluorophore was labeled at H4 E63C. During imaging, continuous 642-nm illumination (∼2 kW/cm^2^) was used to excite Alexa Fluor 647 molecules and switch them into a dark state, and continuous 405-nm illumination (0-1 W/cm^2^) was applied to stochastically reactivate fluorophores to the emissive state. The laser power was adjusted to ensure that only a sparse, optically resolvable subset of fluorophores were active at any given time. Movies consisting of 40,000 image frames were acquired at 100 Hz with a pixel size of 160 nm. Reconstructed 3D STORM images were generated using either every 1,000 or all 40,000 frames.

## Results

### Nucleosome arrays form liquid-like droplet condensates at 150 mM NaCl

Nucleosome arrays were reconstituted with human histone octamers and a DNA template containing 12 repeats of the Widom 601 nucleosome positioning sequence with a 25 bp linker length (Fig. S1). Saturation of reconstituted nucleosome arrays was confirmed by native gel electrophoresis, transmission electron microscopy (TEM), and analytical ultracentrifugation (Figs. S2 – S4). These nucleosome arrays at 200 nM undergo liquid-liquid phase separation (LLPS) into droplet condensates when the ionic environment is set at 150 mM NaCl (Fig. 1). This result further confirms the proper assembly of nucleosome arrays. A previous report has shown that the addition of various physiological monovalent salts (e.g., 150 mM NaCl) to 12-mer nucleosome arrays at a similar level of concentration results in liquid-like droplet condensates^27^. Fusion of droplets on a few seconds timescale confirms the liquid-like nature of these condensates (Fig. 1b). Using fluorescently labeled histone H4 at E63C with Alexa Fluor 647, we found that the concentrations of the nucleosomes in the chromatin droplets are 326 ± 26 μM, in excellent agreement with a previous report (Fig. S5)^27^. The fast recovery of fluorescence after photobleaching (Figs. 1c and S6) also confirms the efficient and constant motions of nucleosomes inside the condensed droplets. The condensates remain liquid-like at least for 2 hours after adding 150 mM NaCl.

**Fig. 1:**
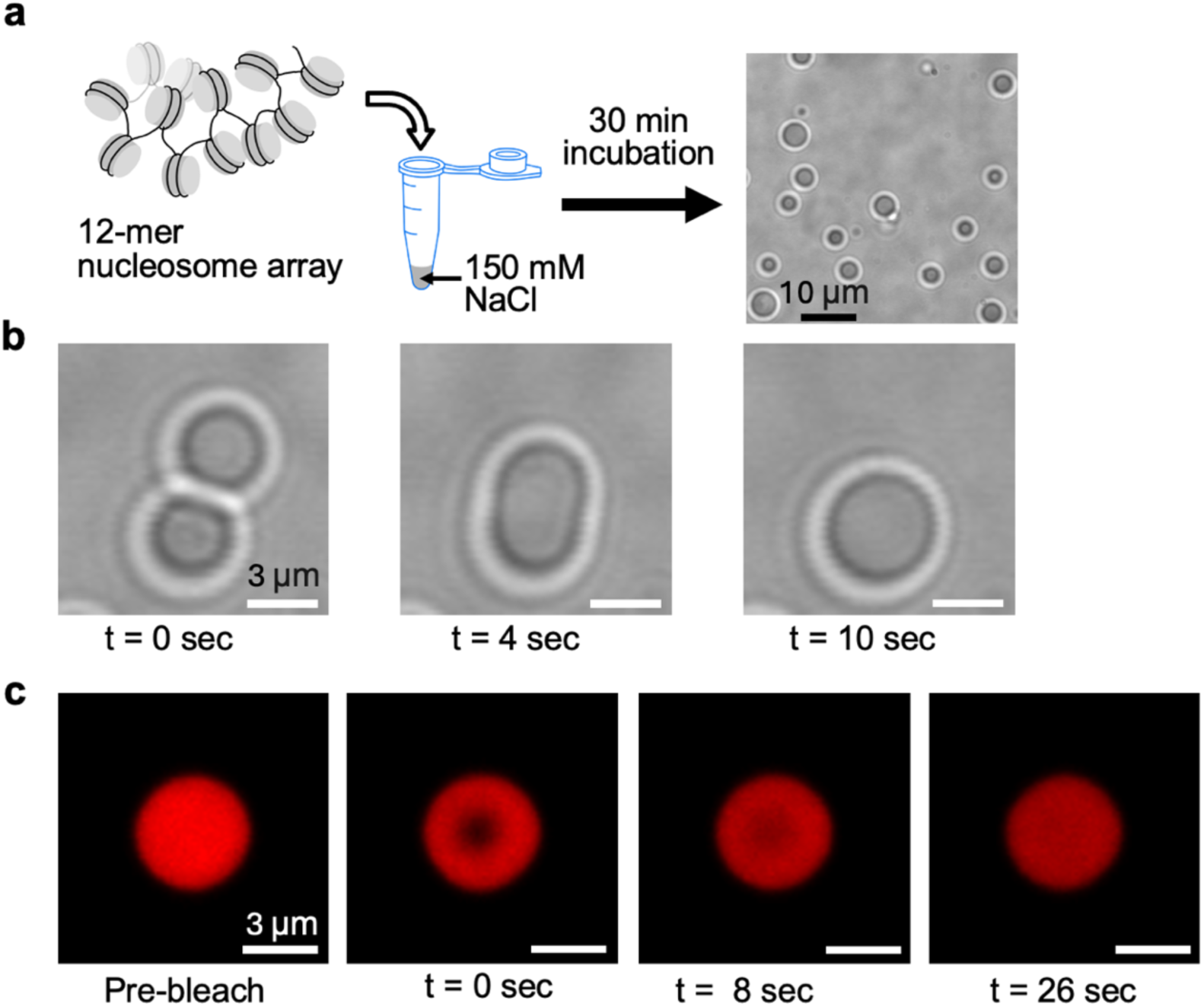
Formation of liquid-like condensates by nucleosome arrays at a physiological salt concentration of 150 mM NaCl. **a** Experimental scheme to form chromatin condensates: Dodecamer nucleosome arrays at 200 nM incubated in a buffer containing 150 mM NaCl result in condensate droplets. The scale bar is 10 μm. **b** Droplets fuse with each other in a few seconds timescale, confirming their liquid-like nature. The scale bars are 3μm. **c** Fluorescence recovery after photobleaching (FRAP) also confirms the liquid-like nature of the droplets. The scale bars are 3μm.

*The lysine residues on histone H4 N-terminal tail are the main drivers of chromatin LLPS* Histone acetylations have been coupled to the weakening of phase separation of nucleosomes and nucleosome arrays^27,36^. In order to investigate the roles of H3 and H4 tails and their acetylation states in regulating LLPS of nucleosome arrays, we prepared four types of nucleosome array samples containing a modified histone core with tailless H3 (gH3), tailless H4 (gH4), an H3 tail acetylation mimic (H3KQ), and an H4 tail acetylation mimic (H4KQ). In H3KQ and H4KQ, the lysine residues in the N-terminal tail (K4, K9, K14, K18, K23, and K27 of H3 and K5, K8, K12, K16, and K20 of H4) are mutated to glutamine to mimic acetylation^51–54^. Among these four samples, only the H3KQ arrays result in chromatin droplets (Fig. 2a). The H4KQ arrays show no sign of droplet formation up to 500 nM array concentration and 200 mM NaCl concentration for 2 hours, while the gH3 and gH4 arrays result in irregularly shaped aggregates (Fig. 2a). These results suggest that H3 and H4 tails play distinct roles in chromatin phase separation and that the H4 tail lysine residues are the main drivers of chromatin LLPS.

**Fig. 2:**
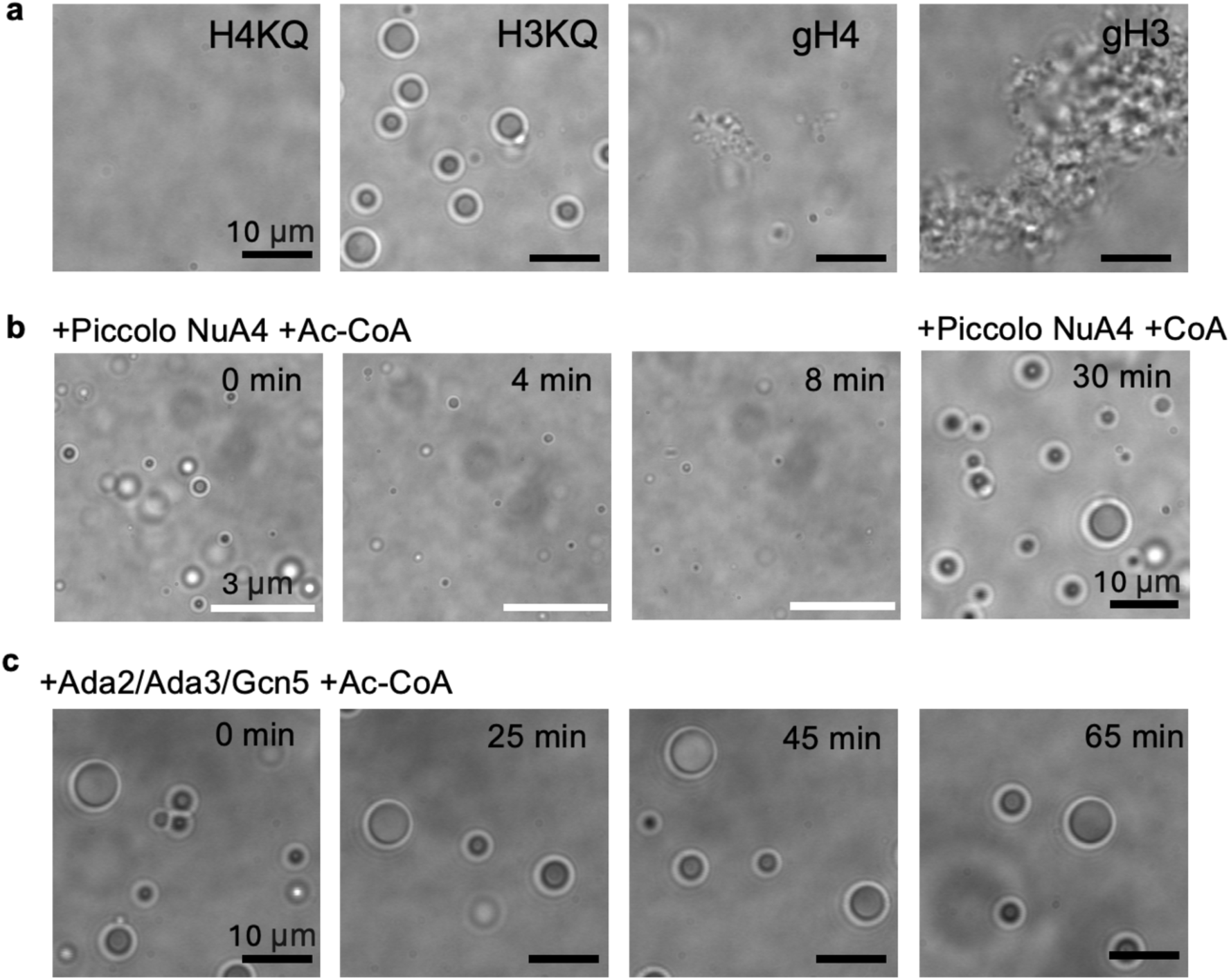
Roles of Histone H4 and H3 tails in regulating phase separation of nucleosome arrays. **a** Nucleosome arrays with the H4 tail acetylation mimic (H4KQ) do not form condensates, but those with the H3 tail acetylation mimic (H3KQ) form spherical phase-separated droplets. Nucleosome arrays with H4 or H3 tails truncated (gH4 or gH3) form solid or gel-like aggregates. The scale bars are 10 μm. **b** *In situ* H4 tail acetylation by Piccolo NuA4 histone acetyltransferase dissolves the phase-separated droplets already formed with nucleosome arrays with no modifications, confirming the effect shown in **a**. Incubation with the enzyme and coenzyme A (CoA) instead of acetyl-CoA (Ac-CoA) does not dissolve the condensates, confirming that the effect is acetylation-dependent. The white scale bars are 3 μm. **c** *In situ* H3 tail acetylation with Ada2/Ada3/Gcn5 histone acetyltransferase does not affect phase separation of nucleosome arrays. The scale bars are 10 μm.

To further confirm the effect of H4KQ, we tested if H4 tail acetylation with histone acetyltransferase (HAT) also inhibits LLPS at a physiological nucleosome concentration. We employed the Piccolo NuA4 complex that acetylates histone H4 tail lysine residues^55,56^. Upon adding Piccolo NuA4 to chromatin droplets formed with unmodified (WT) nucleosome arrays, we observed the gradual dissolution of the droplets (Fig. 2b). The fast enzyme access to histone tails further validates that droplets are liquid-like and can freely exchange materials with the outer environment. Incubation with Piccolo NuA4 and coenzyme-A (Co-A) instead of acetyl Co-A (Ac-CoA) induces no difference in the droplets, confirming that the change is acetylation-dependent (Fig. 2b). Upon a closer look at the droplets, we observed crumbling from the inside, loss of the round outer fringe, and shrinkage of the droplet size within 10 – 15 minutes. We also tested H3 tail acetylation *in situ* with the Ada2/Ada3/Gcn5 complex^57,58^. As expected, no changes were observed in droplet formation, further validating the observation with the H3KQ acetylation mimic. These results confirm that the H4 tail lysine residues are the main drivers of chromatin LLPS regardless of the nucleosome concentration up to a physiological level of a few hundred μM.

### Histone chaperone Nap1 dissolves solid or gel-like chromatin aggregates

The gH3 and the gH4 array samples result in solid or gel-like condensates that do not fuse (Fig. 2a). In particular, gH3 arrays induce large aggregates, while gH4 arrays do so to a much lesser extent. We hypothesize that the aggregation is caused by nucleosome disassembly upon nucleosome destabilization due to the lack of the H3 tails. Tailless H3 induces nucleosome destabilization although it does not result in spontaneous disassembly of mono-nucleosomes^35,46^. However, nucleosomes destabilized at a mono-nucleosomal level would lead to their disassembly when they are condensed. This is because the transiently exposed histone core facilitated by the destabilized DNA wrapping would be prone to random histone-DNA interactions in a condensed phase, leading to nucleosome disassembly.

To test this hypothesis, we added histone chaperone Nap1 to the aggregates formed with gH3 arrays. We observed that upon the addition of Nap1 at a molar ratio of 1:2 (nucleosome:Nap1), the solid or gel-like aggregates dissolve away within a few minutes (Fig. 3a). When Nap1 is added to gH3 arrays at a molar ratio of 1:2 (nucleosome:Nap1) before inducing LLPS, the aggregate formation was significantly reduced (Fig. 3b). No change in LLPS was observed when Nap1 was added to the condensates formed with WT arrays at a molar ratio up to 1:8 (nucleosome:Nap1) (Fig. 3c). These results validate our hypothesis of disassembled nucleosomes and random histone-DNA mixture in condensed chromatin containing tailless H3. The fast action of Nap1 with the pre-formed aggregates supports that these aggregates are more permeable gel-like than solid.

**Fig. 3:**
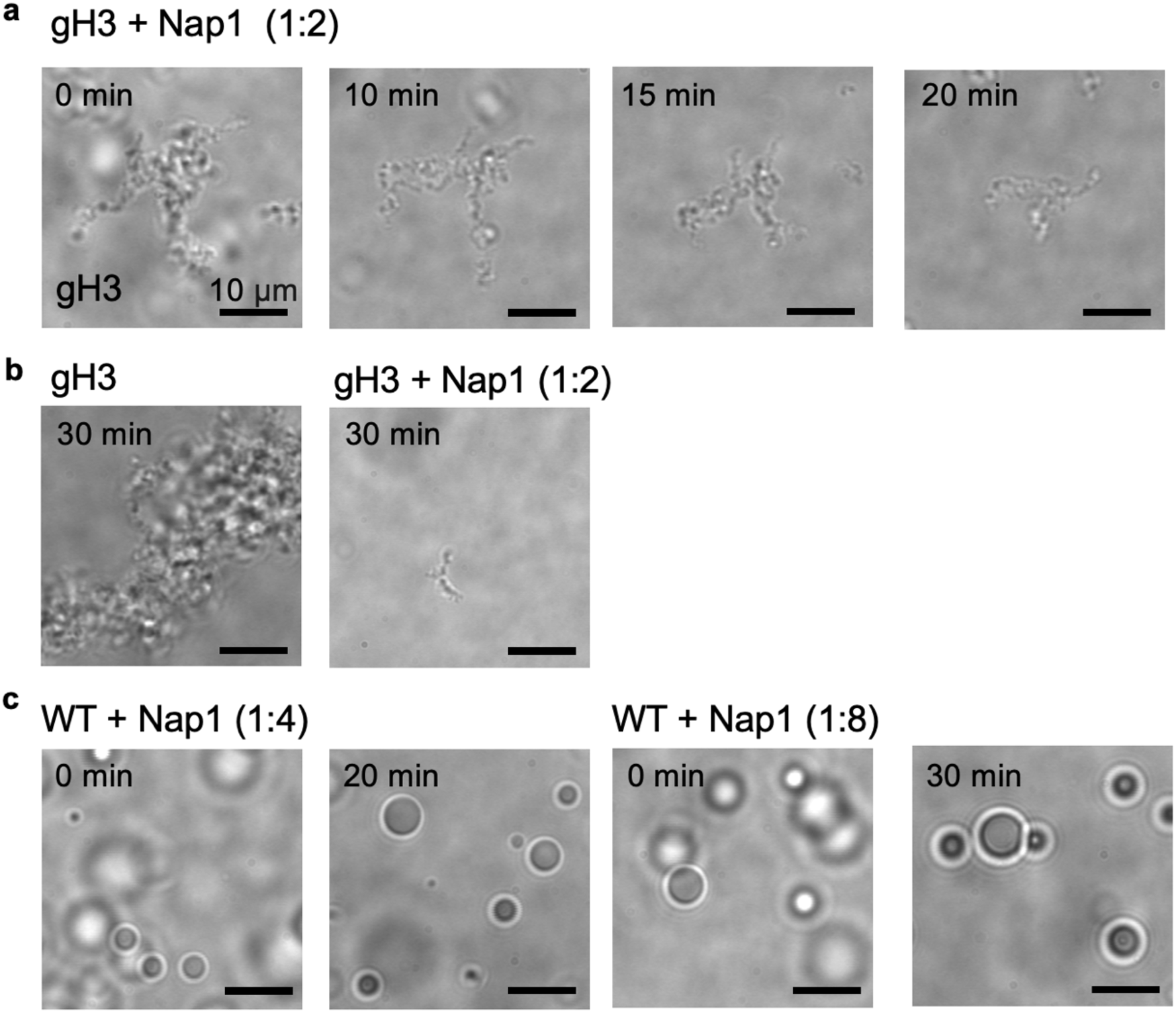
Dissolution of solid or gel-like chromatin aggregates by histone chaperone Nap1. **a** Histone chaperone Nap1 dissolves pre-formed chromatin aggregates with nucleosome arrays containing tailless histone H3 (gH3) at a molar ratio of 1:2 (nucleosome:Nap1). **b** Histone chaperone Nap1 pre-mixed with gH3 nucleosome arrays at a molar ratio of 1:2 (nucleosome:Nap1) before adding 150 mM NaCl significantly inhibits the formation of solid or gel-like chromatin aggregates. **c** The effect of Nap1 presented in **a** is not observed with spherical droplets formed with arrays with no modification (WT) up to a molar ratio of 1:8 (nucleosome:Nap1) for 30 min. All scale bars are 10 μm.

*H3KQ and histone chaperone Nap1 make liquid-like chromatin condensates more fluidic*.

The histone H3 tail lysine residues play important roles in stabilizing nucleosomes by interacting with DNA^30,31,59,60^. Therefore, their acetylation should affect the dynamics of DNA-histone interactions at both the nucleosome and the chromatin levels. To test this hypothesis, we measured the dynamics of fluorescence recovery after photobleaching (FRAP) within the droplets formed with WT and H3KQ arrays. We employed fluorescently labeled histone H4 at E63C (Tet) and H2B at T115C (Di)^61^. We observed at least 2-fold faster recovery on average with H3KQ arrays than with WT arrays in both Tet- and Di-labeled cases (Fig. 4a). The FRAP recovery fractions are comparable in the two cases with no significant difference. These results indicate that nucleosome motions are facilitated in H3KQ droplets, suggesting that DNA-histone dynamics are enhanced at the length and time scales of nucleosome motions. This effect is likely due to weakened interactions between DNA and H3 tail lysine residues. Combined with the results from gH3 arrays that induce nucleosome disassembly in a condensate, these observations suggest that the H3 tail, even when the lysine residues are acetylated and lose their positive charges, still plays a significant role in stabilizing nucleosomes in condensed chromatin.

**Fig. 4:**
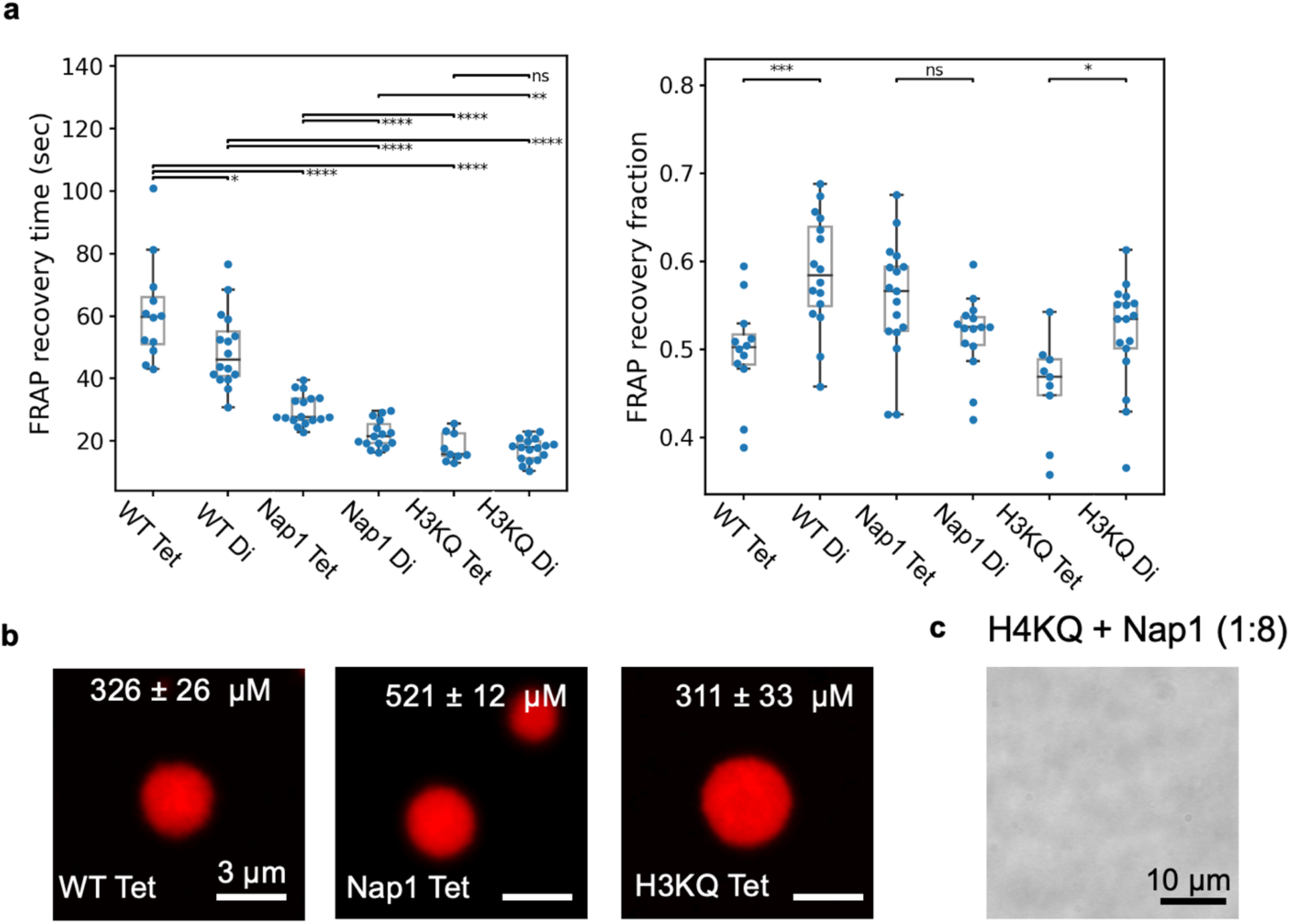
Enhanced nucleosome dynamics in condensed liquid-like droplets by histone chaperone Nap1 and H3 tail acetylation mimic. **a** FRAP recovery times and recovery fractions are plotted for the condensates formed with unmodified nucleosome arrays (WT) labeled at the histone H4 E63C (Tet) and at H2B T115C (Di), unmodified arrays in the presence of Nap1 (Nap1) labeled at Tet and Di, and H3KQ arrays (H3KQ) labeled at Tet and Di. The recovery times of unmodified nucleosome arrays labeled at the histone H4 E63C (Nap1 Tet) and at H2B T115C (Nap1 Di) in the presence of Nap1 are faster than those in its absence, indicating that Nap1 enhances nucleosome dynamics in condensates. The FRAP recovery times of unmodified nucleosome arrays labeled at histone H4 E63C in the presence and absence of Nap1 (WT Tet and Nap1 Tet) are slower than those of unmodified nucleosome arrays labeled at histone H2B T115C (WT Di and Nap1 Di), indicating that histone H2A-H2B dimers diffuse faster than nucleosomes. Nucleosome arrays with H3KQ show dramatically faster FRAP recovery compared to the unmodified arrays (WT). FRAP recovery fractions are in the range of ∼50 - ∼60 % in all cases. A higher FRAP recovery fraction of the Di-labeled unmodified arrays (WT Di) than that of the Tet-labeled WT arrays (WT Tet) is noted. The same trend is noted with H3KQ arrays (H3KQ Tet vs H3KQ Di). The significances shown are from two-sided student’s t-test (*: p ≤0.05, **: p≤0.01, ***: p≤0.001, ****: p≤0.0001, ns: not significant). The sample size n = 12, 16, 17, 15, 9, and 17, respectively for the WT Tet, WT Di, Nap1 Tet, Nap1 Di, H3KQ Tet, and H3KQ Di cases. The sample size refers to the number of distinct droplets tested in each case. The samples and the measurements were made on at least two different days. The values marked on the box plots represent mean ± standard deviation. **b** Concentrations of the nucleosome arrays in the three cases (WT, Nap1, and H3KQ) reveal an elevated concentration of nucleosomes in the presence of Nap1. The concentrations in the WT and H3KQ droplets are the same within error. The scale bars are 3 μm. The sample size n = 15, 18, and 21 respectively for the WT Tet, Nap1 Tet, and H3KQ Tet cases. The sample size refers to the number of distinct droplets tested in each case. The samples and the measurements were made on at least two different days. **c** The effect of Nap1 in elevating the nucleosome concentration in chromatin condensates does not result in droplet formation with H4KQ arrays up to two hours.

To test the effect of Nap1 on regulating DNA-histone dynamics in condensed chromatin, we monitored the FRAP recovery dynamics within the droplets formed with WT arrays in the presence and absence of Nap1 (Fig. 4a). According to the observations, Nap1 makes the chromatin droplets more fluidic in the relevant length and time scales in both Tet- and Di-labeled cases. Spontaneous DNA-histone dynamics on a few milliseconds timescale result in reversible and repetitive partial disassembly and reassembly of nucleosomes. Our results suggest that Nap1 facilitates and stabilizes these microscopic dynamics to induce a macroscopic change in the condensed nucleosome motions on a few to a few tens of seconds timescale^62^.

To examine the mechanism of FRAP recovery, we compared the dynamics from the two different labeling positions which are H4E63C (Tet) and H2BT115C (Di). The Tet-labeled arrays show a slower recovery time than the Di-labeled ones in the WT and the Nap1 cases (Fig. 4a). The difference is not significant in H3KQ arrays, likely because the recovery times are already too short to reliably gauge any differences at the resolution of our measurements. We propose that this difference is due to spontaneous histone H2A-H2B dimer exchange between nucleosomes^43^. The higher fraction of recovery in the Di-labeled droplets further supports this mechanism (Fig. 4a). As for the Nap1 case, the FRAP recovery fractions are not significantly different between the Tet- and Di-labeled arrays. This likely originates from that Nap1-mediated DNA-histone dynamics apply to both (H3-H4)_2_ tetramers and H2A-H2B dimers and that these dynamics contribute significantly to the FRAP behavior. The FRAP recovery times in the Nap1 Tet/Di cases are nearly two-fold faster than those in the WT Tet/Di cases. Regardless, our results strongly support that histone H2A-H2B spontaneously exchanges between intact nucleosomes in a condensed phase. This also suggests a mechanism for spontaneous H2A-H2B diffusion within condensed chromatin, facilitated by Nap1.

### The effects of the H3 acetylation mimic and Nap1 are not due to changes in the nucleosome concentration

We further validated the roles of H3 tail lysine residues and Nap1 in facilitating chromatin dynamics in a condensed phase by showing that the nucleosome concentration within a droplet does not decrease in these cases. We measured the concentrations of the nucleosomes in the three cases (WT, Nap1, and H3KQ) with the Tet-labeled samples (Fig. 4b). We found that the concentrations stayed constant within error between the WT (326 ± 26 μM) and the H3KQ (311 ± 33 μM) cases, confirming that the H3KQ effect on making chromatin more dynamic is not due to decreased nucleosome density in the droplets. Strikingly, we found that the nucleosome concentration increases significantly upon the addition of Nap1 from 326 ± 26 (WT) to 521 ± 12 μM (Nap1). This effect is likely due to the elevated thermodynamic stability of nucleosomes in the presence of Nap1. This function of Nap1 in elevating nucleosome concentration in condensed chromatin has never been reported. To this end, we tested if the droplet formation is induced specifically by the inter-nucleosomal interactions between and within nucleosome arrays that are mediated by histone H4 tail lysine residues, or generally by an increased nucleosome concentration that can be caused by either H4-mediated inter-nucleosomal interactions or various other factors such as Nap1. When Nap1 was added to H4KQ arrays at up to a 1:8 (nucleosome:Nap1) molar ratio, no droplet formation was observed up to two hours. This result further confirms that H4 tail lysine residues are the main drivers of chromatin LLPS, regardless of the nucleosome concentration or stability within a condensate.

### STORM imaging reveals a relatively static structural scaffold in condensed chromatin droplets

The FRAP results (Fig. 4a) indicate that the fluorescence recovery is limited to ∼50 – ∼60 %. This observation led us to a hypothesis that condensed chromatin droplets contain a network of nucleosome arrays that form a relatively immobile structural scaffold of the droplet, moving too slowly to be captured during our FRAP measurements (up to 216 sec). To test this hypothesis, we used stochastic optical reconstruction microscopy (STORM) to image nucleosomes that are relatively static on the time scale of 10 seconds in droplets (Fig. 5). We took fluorescence images of chromatin droplets containing Tet-labeled nucleosome arrays every 10 ms for 400 seconds, resulting in 40,000 fluorescence images. Every 1,000 images, corresponding to a 10-second period, were processed to generate a 10-second “Total” STORM image as shown in Fig. 5 (upper left in WT and H3KQ). The full imaging duration of 400 seconds produced 40 of such 10-second STORM images. Within each 10-second STORM image, we identified “Clusters” by grouping 5 or more histone H4 molecule localizations not deviating more than 110 nm from each other, where the length 110 nm is an approximate contour length of 12-mer nucleosome arrays determined from TEM images. Only retaining the identified clusters and removing the H4 molecule localization outside the clusters in the 10-second STORM image resulted in a 10- second “Clusters” STORM image. Examples of such clusters are shown in Fig. 5, upper right, in WT and H3KQ. Comparing the STORM images of “Total” and “Clusters” in Fig. 5 reveals that many H4 molecules remain relatively static during a 10-second period, while the others are more mobile with a motion speed at or above our 100 Hz image acquisition rate. Forty 10-second STORM images were overlaid to generate “Total” and “Clusters” views, representing all nucleosomes and relatively static nucleosome clusters, respectively, identified over the full 400- second imaging session (Fig. 5, lower left and right, respectively, in WT and H3KQ). From these STORM images, it is evident that the entirety of the droplet interior is full of relatively static nucleosomes on a 10-second timescale represented by the dots on a “Clusters” image. It is also evident that there are mobile nucleosomes with a motion speed at or above our 100Hz image acquisition rate on a 10-second timescale shown as the dots on a “Total” image which do not appear on the corresponding “Clusters” image in both the WT and the H3KQ cases. Overall, STORM imaging confirms that condensed chromatin droplets contain both mobile and relatively immobile scaffolding nucleosomes on a 10-second timescale.

**Fig. 5:**
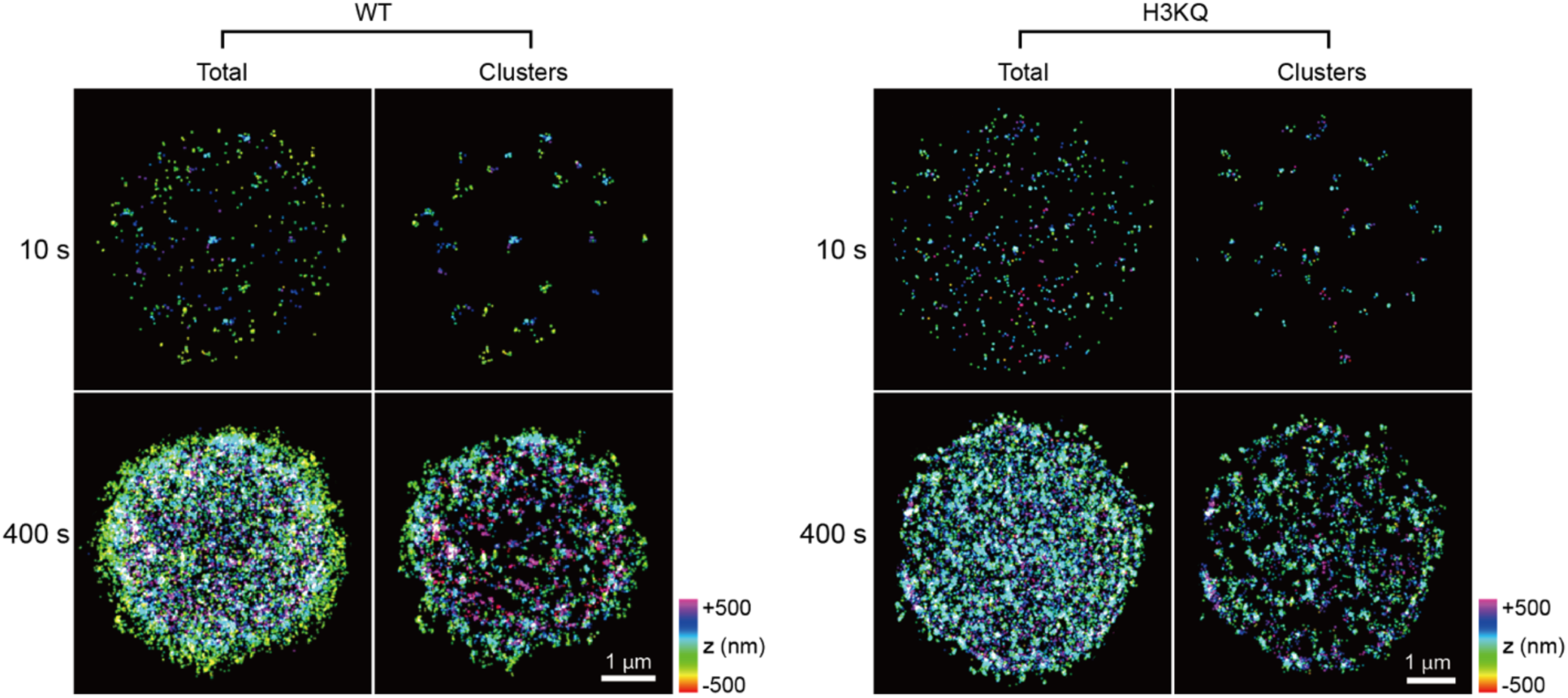
Optical super-resolution images of liquid-like chromatin droplets. Fluorescently labeled nucleosome arrays at H4 E63C were imaged with STORM. The “Total” images for 10 s represent the nucleosome particles identified during a 10-sec period of STORM imaging. The “Clusters” images for 10 s represent five or more histone H4 molecules clustered together within 110 nm from each other during a 10-sec period. The “Total” images for 400 s represent the overlap of the 40 consecutive 10-s “Total” images. The “Clusters” images for 400 s represent the overlap of the 40 consecutive 10-s “Clusters” images. These STORM images of both WT and H3KQ arrays confirm a structural scaffold of nucleosome arrays that are relatively immobile on the timescale of 10 seconds. The z-position of the localized nucleosomes are color coded as shown in the color scale.

### Microrheology confirms low viscosity of droplets in the presence of Nap1 or with H3KQ modification

FRAP recovery times provide insights into how dynamically nucleosomes move within a droplet. FRAP recovery within a droplet reflects the motions of nucleosomes from the mobile fraction of the condensate, which pass through both mobile and immobile nucleosomes and arrays. The motions encounter friction due to collisions most strongly with other nucleosome arrays. Such friction results in slower diffusion, which can be interpreted as a higher viscosity of the diffusion medium. The FRAP recovery times are a convolution of multiple parameters affecting diffusion, such as the viscosity of the solvent and various interactions of the diffusing body with other components in the diffusion medium (e.g., collisions, transient association, and dissociation). The Stokes-Einstein relation between diffusion and viscosity does not accurately reflect such complex dynamics. Moreover, polymeric materials often display viscoelastic behavior that depends on the rate of motion. Therefore, a simple convolution of these factors into FRAP recovery times would not properly capture the nucleosome dynamics and their changes due to modified histones and the presence of Nap1. To this end, we measured the viscoelasticity of chromatin droplets using an optical tweezers setup. We brought a polystyrene bead (1 μm diameter) into a droplet (>4 μm diameter) and sinusoidally oscillated the bead with a ±0.5 μm amplitude (Fig. 6a). We employed a Burgers model, a simple linear viscoelasticity model, with two Maxwell components to interpret the data^49^. The drag and the attenuation in the bead motion at various oscillation frequencies contain all the relevant pieces of information (Fig. 6a). The storage and loss moduli at each frequency can be extracted from these bead motions. Fitting the frequency dependence of the moduli results in the relaxation times and the viscosities (Fig. 6b). A single Maxwell component did not fit the data. The two-component fitting worked well, and the results are shown in Fig. 6c. The faster relaxation component has a relaxation time of a few ms which is 2-3 orders of magnitude shorter than the slower component, which shows a relaxation time of a few hundred ms to a few seconds. The viscosity of the faster component is 2- 3 orders of magnitude lower than that of the slower component and only 2-3 times higher than that of water (0.001 Pa⋅s at 25 °C). Therefore, the faster component is likely due to the dynamics of the mobile arrays freely diffusing within a droplet. The faster component should result in diffusion resistance from the mobile arrays, while the slower component exerts resistance from the structural scaffold. The faster relaxation would be governed mainly by intra-array conformational dynamics of mobile arrays during diffusion while the slower relaxation would be governed mainly by rearrangements of the intra- and inter-array networks in the scaffold. The 2- 3 orders of magnitude difference in their timescales is well aligned with that the network reconfiguration of the scaffold arrays should be much slower than the conformational changes of mobile arrays^63^. Under the condition of 2-3 orders of magnitude difference in the viscosity, the network reconfiguration time is approximately the relaxation time of the slower component which is a few hundred ms to a few seconds. Processes that are faster than this timescale will experience both viscous and elastic properties of the chromatin while slower processes will experience mostly the viscous nature with the viscosity values shown in Fig. 6c.

**Fig. 6:**
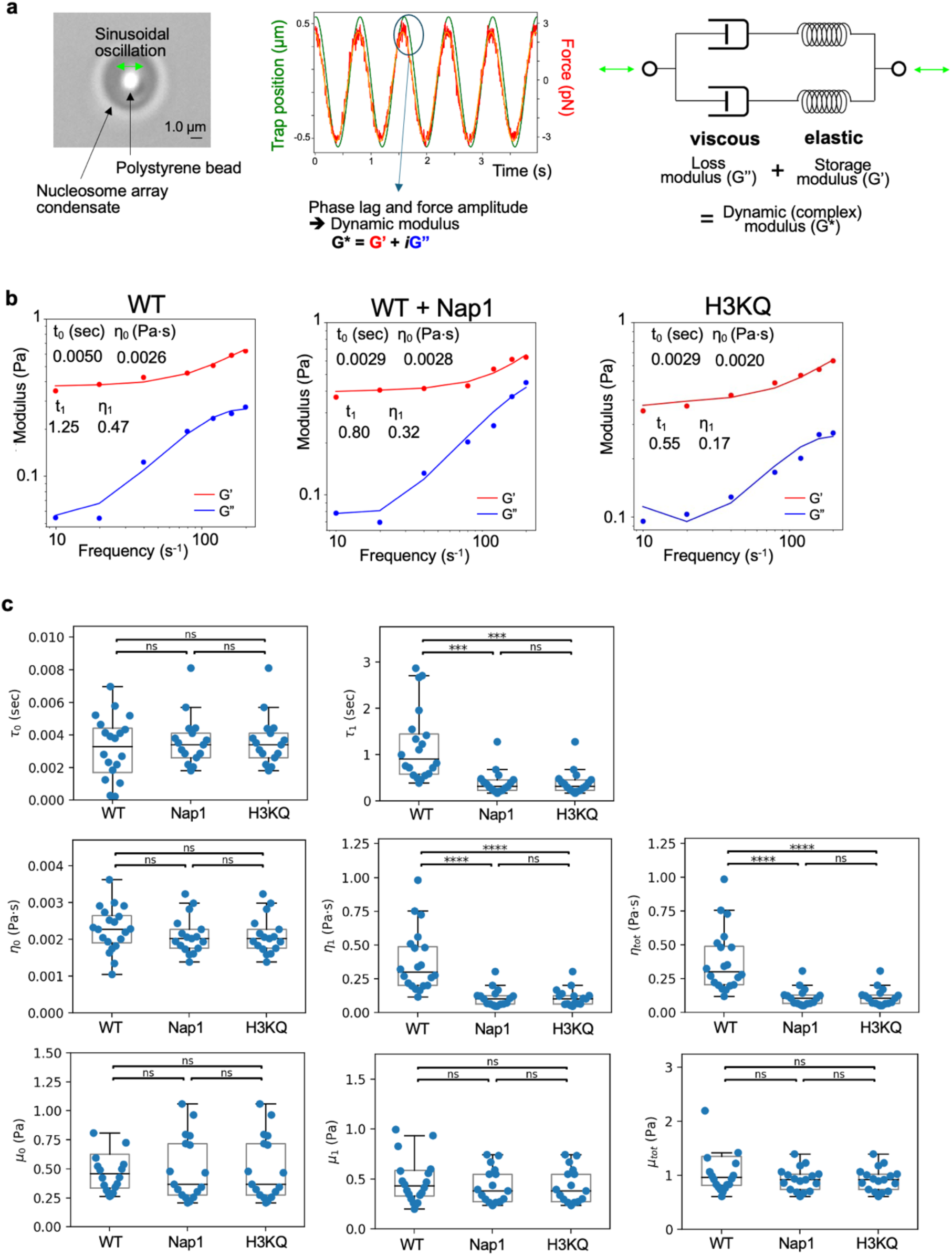
Microrheology measurements of condensed droplets and the effects of histone chaperone Nap1 and H3KQ acetylation mimic. **a** Experiments were set up based on optical tweezers to measure the dynamic modulus of nucleosome array droplets. A 1 μm polystyrene bead is brought inside a condensate whose diameter is at least 4 μm. The bead is sinusoidally oscillated at 7 different frequencies (left). The oscillation amplitude of the bead and the phase lag between the bead and the trapping beam are monitored (center). The lag and the attenuated amplitude of the bead are fit with a Burgers model with two Maxwell components (right). **b** Examples of dynamic modulus fitting and results for unmodified arrays (WT), WT arrays with Nap1 (Nap1), and arrays with the H3 tail acetylation mimic (H3KQ). **c** The fitting results are summarized for the two relaxation time components τ_0_ and τ_1_, the two viscosity components η_0_ and η_1_, and their sum η_tot_, the two corresponding elastic modulus μ_0_ and μ_1_, and their sum μ_tot_ . The longer time relaxation component and its viscosity is significantly shorter and lower in the Nap1 and H3KQ cases than the WT case. As the longer relaxation component is the dominant component, the total viscosity is also significantly lower with the Nap1 and H3KQ cases, revealing less viscous and more fluidic condensates induced by these changes. The sample size n = 19, 20, and 16, respectively for the WT, Nap1, and H3KQ cases. The sample size refers to the number of distinct droplets tested in each case. The samples and the measurements were made on at least two different days. The values marked on the box plots represent mean ± standard deviation. The significances shown are from two-sided student’s t-test (*: p ≤0.05, **: p≤0.01, ***: p≤0.001, ****: p≤0.0001, ns: not significant).

According to the analysis, the relaxation time and the viscosity of the faster relaxation component stay constant within error in the WT and H3KQ cases. This may be because H3KQ would exert two opposing effects on the intra-array conformational dynamics of mobile arrays. One is to make the mobile arrays more dynamic due to intra-array DNA-histone dynamics facilitated by H3KQ. From the point of mobile arrays casting barriers to the motions of another body, this effect will make the bead motion more efficient, consequently resulting in a faster relaxation response and a lower viscosity. The other effect is from the perspective of the array motions themselves. The effect described above due to H3KQ will result in a larger hydrodynamic radius, thereby impeding the diffusion of mobile arrays. Such reduced diffusion would result in a slower relaxation response and a higher viscosity. Therefore, these two opposing effects will cancel each other, and the relaxation time and viscosity may stay apparently unchanged. This explanation can also account for the unchanged relaxation time observed in the presence of Nap1 although the viscosity is elevated likely due to a higher concentration of nucleosome arrays induced by Nap1.

On the other hand, the slower relaxation component shows significantly shorter relaxation time and lower viscosity with Nap1 and H3KQ arrays than with WT arrays. These results suggest that the intra- and inter-array nucleosome dynamics in the structural scaffold take place on the order of a few hundred ms to a few seconds and that the network reconfiguration dynamics is significantly facilitated by Nap1 or H3KQ. Therefore, the faster FRAP recovery induced by Nap1 and H3KQ must be largely due to facilitated network reconfiguration dynamics of the structural scaffold in a condensate, which originates from eased intra- and inter-array nucleosome motions. These results support that the viscous resistance of condensed chromatin can be lessened by histone chaperone and histone H3 tail acetylation, thereby facilitating the structural fluidity of intercalated nucleosome arrays at a chromatin level.

## Discussions

Nucleosome arrays are naturally prone to forming condensates via LLPS as they contain both unstructured positively charged protein chains and negatively charged DNA which are the main constituents of various LLPS systems^27,29,64^. Our results indicate that these condensed droplets reach nucleosome concentrations comparable to those found in the cell nucleus, serving as an excellent platform for investigating various chromatin components in a condensed phase. In condensed chromatin droplets, the dynamics of DNA-histone interactions and their regulations are not straightforward to model as nucleosomes are closely packed to each other while their motions are constrained by being linked in an array. One such example is spontaneous histone H2A-H2B exchange between nucleosomes. When nucleosomes freely diffuse in solution, they collide with each other and exchange H2A-H2B when the collision coincides with spontaneous partial unwrapping of nucleosomal DNA^65^. The timescale of spontaneous H2A-H2B exchange between freely diffusing nucleosomes at a few hundred μM concentration is on the order of a few seconds per nucleosome. However, it is hard to predict if such exchange is feasible and significant in a condensed phase because nucleosome motions are constrained. Our results show significantly facilitated diffusion dynamics of the nucleosomes labeled at H2A-H2B over those labeled at (H3-H4)_2_ in the WT and Nap1 cases. These results support spontaneous H2A-H2B exchange between nucleosomes within the structural scaffold of a condensate. This exchange would constitute a diffusion mechanism of H2A-H2B in condensed chromatin. The diffusion of H2A-H2B within condensed chromatin may provide an efficient way to remove their variants and post-translational modifications once their functions are fulfilled. Further investigation and quantitative analysis of the dynamics of modified H2A–H2B, including their epigenetic marks and variants, will be crucial for uncovering their biophysical mechanisms of gene regulation.

Acetylation of the lysine residues on the N-terminal tail of histone H4 have been associated mostly with inhibited inter-nucleosomal interactions^60,66^. The effect is due to the modulated H4 interactions with DNA and the acidic patch of the histone core across nucleosomes. It has been suggested that chromatin phase separation is induced by both short- and long-range inter-nucleosomal interactions between DNA and histone^67,68^. The short- and long-range interactions would be equivalent to intra- and inter-array interactions between nucleosomes in our system^69^. Both of these interactions would be inhibited by H4 tail acetylation. Our results indicate that such inhibition is sufficient to block chromatin phase separation of nucleosome arrays. The N-terminal tail of histone H3 also plays a role in inter-nucleosomal interactions, but to a lesser extent than that of H4, which is supported by efficient droplet formation with H3KQ arrays and large aggregate formation with gH3 arrays. Therefore, our results suggest that H4-tail acetylation would exert a direct and strong impact on chromatin condensation and its accessibility. Deacetylations of H4 tail lysine residues have been implicated in chromatin condensation during mitosis and their acetylation is inversely correlated with inhibition of chromatin condensation in prophase.^70,71^

Acetylation of the lysine residues on the N-terminal tail of histone H3 has been coupled to destabilized nucleosomes and elevated DNA accessibility within nucleosomes^30,31,59,60^. The effects are largely due to their weakened interactions with DNA. Such destabilization of the nucleosome structure is not to the extent where nucleosomes will disassemble spontaneously. Rather, it eases mechanical unwrapping of the outer region of nucleosomes, enhances thermal and salt-induced disassembly, and facilitates transcription by inhibiting an entry pause to the nucleosome^34,35,46^. These changes are correlated with making nucleosomes more flexible on the timescales of these events which are much longer than that of the spontaneous “breathing” motions of the nucleosome at milliseconds^62^. Such elevated flexibility would be observable as more dynamic motions of nucleosomes at both the nucleosome and the array levels. Our FRAP and microrheology results provide direct supports to these changes by showing enhanced diffusion and faster relaxation on a few hundred ms to a few tens of seconds timescale. Our results also revealed that the facilitated motions and fluidity of nucleosomes and nucleosome arrays induced by H3KQ are not sufficient to inhibit chromatin phase separation. Therefore, H3 tail acetylation would mainly function to enhance DNA-histone dynamics and protein diffusion within condensed chromatin.

Unlike gH4 arrays, gH3 arrays result in large-scale gel-like aggregates. As both tails are important in stabilizing DNA-histone interactions within and across nucleosomes, we suggest that these aggregates are seeded by disassembled nucleosomes upon their initial condensation at 150 mM NaCl. Nucleosome core particles containing gH3 or gH4 have nearly unchanged structures compared with canonical nucleosomes^72,73^. However, their dynamics and structural stabilities should be altered, which may result in the complete disassembly of nucleosomes upon closer and sustained contact with each other. Such contacts would involve interactions between transiently exposed histones and transiently unwrapped DNA in an intra- and inter-nucleosomal way, leading to nucleosome disassembly and aggregation of random DNA-histone mixtures.

Histone chaperone Nap1 mediates these unstable DNA-histone interactions, dissociates histones from DNA, and shuttles them back to DNA for proper nucleosome assembly^40^. Our result showing Nap1-induced dissolution of gel-like aggregates strongly suggests that the aggregates are permeable gel-like and that they are formed by random interactions between DNA and histone. Therefore, histone H3 and H4 tail residues other than the lysine residues must play critical roles in stabilizing nucleosomes in condensed chromatin.

Faster FRAP recovery with Nap1 further confirms its role in dynamically mediating DNA-histone interactions. Such mediation has been reported from the thermodynamic perspectives^40,74^. However, it remained unknown how fast Nap1 mediates DNA-histone interactions to induce an observable change at a macroscopic level. Our FRAP results indicate that Nap1 facilitates DNA-histone dynamics within and across intact nucleosomes on the order of a few to a few tens of seconds that is the diffusion timescales of nucleosomes in our chromatin condensates. This is a surprising point of Nap1’s function in enhancing DNA-histone dynamics on this short timescale. As FRAP measurements report diffusion within and of the mobile fraction of the condensed droplet, they may not directly capture the properties of the immobile fraction which is important in chromatin condensation *in vivo*. The existence of the mobile and relatively immobile fractions are confirmed with the ∼50 – ∼60 % FRAP recovery and the STORM images. To this end, we investigated if the effect of Nap1 on enhancing DNA-histone dynamics applies to the relatively immobile structural scaffold with microrheology measurements. Unlike FRAP, these measurements report the properties of both the mobile and the immobile fractions of a droplet. Our results show that Nap1 enhances DNA-histone dynamics during network reconfiguration in the relatively immobile fraction on a relaxation timescale of a few hundred ms to a few seconds. This timescale is relevant to the diffusion and binding of various enzymes in chromatin such as RNA polymerase II and chromatin remodelers, and their actions in reading and altering chromatin^75^. Therefore, Nap1 can function as an effective regulator of these enzymes on the timescale relevant to their functions in condensed chromatin without directly interacting with them.

Our microrheology measurements revealed two relaxation components. The faster relaxation component is likely due to intra-array conformational dynamics that would be relevant to short-range inter-nucleosomal interactions in condensed chromatin, while the slower relaxation component would be due to long-range inter-nucleosomal interactions. Therefore, the slower component would likely mimic a relatively immobile structural scaffold of condensed chromatin formed with long nucleosome arrays *in vivo*. Moreover, the markedly elevated chromatin fluidity observed with Nap1 addition and H3KQ modification are due to the facilitated reconfiguration of the structural scaffold, suggesting broader relevance to the regulation of condensed chromatin structure and dynamics.

## Acknowledgements

This research was supported by NIH grants R35GM148208 to T.L., R35GM142973 to R.Z., and R35GM127034 to S.T. The analytical ultracentrifuge used for this research was funded by NIH (S10OD032215).

## References

1. Brangwynne, C. P., Mitchison, T. J. & Hyman, A. A. Active liquid-like behavior of nucleoli determines their size and shape in *Xenopus laevis* oocytes. Proc. Natl. Acad. Sci. 108, 4334–4339 (2011).

2. Feric, M. et al. Coexisting Liquid Phases Underlie Nucleolar Subcompartments. Cell 165, 1686–1697 (2016).

3. Finn, E. H. & Misteli, T. Molecular basis and biological function of variability in spatial genome organization. Science 365, eaaw9498 (2019).

4. Luger, K., Mäder, A. W., Richmond, R. K., Sargent, D. F. & Richmond, T. J. Crystal structure of the nucleosome core particle at 2.8 Å resolution. Nature 389, 251–260 (1997).

5. Hihara, S. et al. Local Nucleosome Dynamics Facilitate Chromatin Accessibility in Living Mammalian Cells. Cell Rep. 2, 1645–1656 (2012).

6. Kornberg, R. D. & Lorch, Y. Primary Role of the Nucleosome. Mol. Cell 79, 371–375 (2020).

7. Lai, W. K. M. & Pugh, B. F. Understanding nucleosome dynamics and their links to gene expression and DNA replication. Nat. Rev. Mol. Cell Biol. 18, 548–562 (2017).

8. Ahmad, K. & Henikoff, S. Epigenetic Consequences of Nucleosome Dynamics. Cell 111, 281–284 (2002).

9. Berger, S. L. Histone modifications in transcriptional regulation. Curr. Opin. Genet. Dev. 12, 142–148 (2002).

10. Jenuwein, T. & Allis, C. D. Translating the Histone Code. Science 293, 1074–1080 (2001).

11. Lawrence, M., Daujat, S. & Schneider, R. Lateral Thinking: How Histone Modifications Regulate Gene Expression. Trends Genet. 32, 42–56 (2016).

12. Clapier, C. R., Iwasa, J., Cairns, B. R. & Peterson, C. L. Mechanisms of action and regulation of ATP-dependent chromatin-remodelling complexes. Nat. Rev. Mol. Cell Biol. 18, 407–422 (2017).

13. Hammond, C. M., Strømme, C. B., Huang, H., Patel, D. J. & Groth, A. Histone chaperone networks shaping chromatin function. Nat. Rev. Mol. Cell Biol. 18, 141–158 (2017).

14. Reyes, A. A., Marcum, R. D. & He, Y. Structure and Function of Chromatin Remodelers. J. Mol. Biol. 433, 166929 (2021).

15. Avvakumov, N., Nourani, A. & Côté, J. Histone Chaperones: Modulators of Chromatin Marks. Mol. Cell 41, 502–514 (2011).

16. Bannister, A. J. & Kouzarides, T. Regulation of chromatin by histone modifications. Cell Res. 21, 381–395 (2011).

17. Kimura, H. Histone modifications for human epigenome analysis. J. Hum. Genet. 58, 439– 445 (2013).

18. Lee, T.-H. Physical Chemistry of Epigenetics: Single-Molecule Investigations. J. Phys. Chem. B 123, 8351–8362 (2019).

19. Kuryan, B. G. et al. Histone density is maintained during transcription mediated by the chromatin remodeler RSC and histone chaperone NAP1 in vitro. Proc. Natl. Acad. Sci. 109, 1931–1936 (2012).

20. Aguilar-Gurrieri, C., et al. Structural evidence for Nap1-dependent H2A–H2B deposition and nucleosome assembly. EMBO J. 35, 1465–1482 (2016).

21. Tachiwana, H. et al. Nap1 regulates proper CENP-B binding to nucleosomes. Nucleic Acids Res. 41, 2869–2880 (2013).

22. Reinkemeier, C. D. & Lemke, E. A. Raising the ribosomal repertoire. Nat. Chem. 12, 503– 504 (2020).

23. Harada, B. T. et al. Stepwise nucleosome translocation by RSC remodeling complexes. eLife 5, e10051 (2016).

24. Spangler, C. J. et al. DOT1L activity in leukemia cells requires interaction with ubiquitylated H2B that promotes productive nucleosome binding. Cell Rep. 38, 110369 (2022).

25. Carcamo, C. C. et al. ATP binding facilitates target search of SWR1 chromatin remodeler by promoting one-dimensional diffusion on DNA. eLife 11, e77352 (2022).

26. Bernier, M. et al. Linker histone H1 and H3K56 acetylation are antagonistic regulators of nucleosome dynamics. Nat. Commun. 6, 10152 (2015).

27. Gibson, B. A. et al. Organization of Chromatin by Intrinsic and Regulated Phase Separation. Cell 179, 470–484.e21 (2019).

28. Gibson, B. A., et al. In diverse conditions, intrinsic chromatin condensates have liquid-like material properties. Proc. Natl. Acad. Sci. 120, e2218085120 (2023).

29. Strickfaden, H. et al. Condensed Chromatin Behaves like a Solid on the Mesoscale In Vitro and in Living Cells. Cell 183, 1772–1784.e13 (2020).

30. Ausio, J. & Van Holde, K. E. Histone hyperacetylation: its effects on nucleosome conformation and stability. Biochemistry 25, 1421–1428 (1986).

31. Norton, V. G., Imai, B. S., Yau, P. & Bradbury, E. M. Histone acetylation reduces nucleosome core particle linking number change. Cell 57, 449–457 (1989).

32. Kan, P.-Y., Caterino, T. L. & Hayes, J. J. The H4 Tail Domain Participates in Intra- and Internucleosome Interactions with Protein and DNA during Folding and Oligomerization of Nucleosome Arrays. Mol. Cell. Biol. 29, 538–546 (2009).

33. Ghoneim, M., Fuchs, H. A. & Musselman, C. A. Histone Tail Conformations: A Fuzzy Affair with DNA. Trends Biochem. Sci. 46, 564–578 (2021).

34. Kim, T. H., et al. Correlating histone acetylation with nucleosome core particle dynamics and function. Proc. Natl. Acad. Sci. 120, e2301063120 (2023).

35. Brower-Toland, B. et al. Specific Contributions of Histone Tails and their Acetylation to the Mechanical Stability of Nucleosomes. J. Mol. Biol. 346, 135–146 (2005).

36. Hammonds, E. F. et al. Histone H3 and H4 tails play an important role in nucleosome phase separation. Biophys. Chem. 283, 106767 (2022).

37. Park, Y.-J. & Luger, K. The structure of nucleosome assembly protein 1. Proc. Natl. Acad. Sci. 103, 1248–1253 (2006).

38. Bowman, A. et al. The Histone Chaperones Nap1 and Vps75 Bind Histones H3 and H4 in a Tetrameric Conformation. Mol. Cell 41, 398–408 (2011).

39. D’Arcy, S. et al. Chaperone Nap1 Shields Histone Surfaces Used in a Nucleosome and Can Put H2A-H2B in an Unconventional Tetrameric Form. Mol. Cell 51, 662–677 (2013).

40. Andrews, A. J., Chen, X., Zevin, A., Stargell, L. A. & Luger, K. The Histone Chaperone Nap1 Promotes Nucleosome Assembly by Eliminating Nonnucleosomal Histone DNA Interactions. Mol. Cell 37, 834–842 (2010).

41. Correll, S. J., Schubert, M. H. & Grigoryev, S. A. Short nucleosome repeats impose rotational modulations on chromatin fibre folding: Rotational modulations of chromatin folding. EMBO J. 31, 2416–2426 (2012).

42. Muthurajan, U., et al. In Vitro Chromatin Assembly: Strategies and Quality Control. In Methods Enzymol. 573 3–41 (2016).

43. Das, S. K., Huynh, M. T. & Lee, T.-H. Spontaneous histone exchange between nucleosomes. J. Biol. Chem. 299, 105037 (2023).

44. Huang, J. & Tan, S. Piccolo NuA4-Catalyzed Acetylation of Nucleosomal Histones: Critical Roles of an Esa1 Tudor/Chromo Barrel Loop and an Epl1 Enhancer of Polycomb A (EPcA) Basic Region. Mol. Cell. Biol. 33, 159–169 (2013).

45. Barrios, A. et al. Expression and purification of recombinant yeast Ada2/Ada3/Gcn5 and Piccolo NuA4 histone acetyltransferase complexes. Methods 41, 271–277 (2007).

46. Oishi, T. et al. Contributions of histone tail clipping and acetylation in nucleosome transcription by RNA polymerase II. Nucleic Acids Res. 51, 10364–10374 (2023).

47. Das, S. K. et al. Methods to investigate nucleosome structure and dynamics with single-molecule FRET. Methods 215, 17–27 (2023).

48. Alshareedah, I. et al. Sequence-specific interactions determine viscoelasticity and ageing dynamics of protein condensates. Nat. Phys. 20, 1482–1491 (2024).

49. Ghosh, A., Kota, D. & Zhou, H.-X. Shear relaxation governs fusion dynamics of biomolecular condensates. Nat. Commun. 12, 5995 (2021).

50. Huang, B., Wang, W., Bates, M. & Zhuang, X. Three-Dimensional Super-Resolution Imaging by Stochastic Optical Reconstruction Microscopy. Science 319, 810–813 (2008).

51. Dewing, S. M., Phan, T. M., Kraft, E. J., Mittal, J. & Showalter, S. A. Acetylation-Dependent Compaction of the Histone H4 Tail Ensemble. J. Phys. Chem. B 128, 10636–10649 (2024).

52. Szerlong, H. J., Prenni, J. E., Nyborg, J. K. & Hansen, J. C. Activator-dependent p300 Acetylation of Chromatin in Vitro. J. Biol. Chem. 285, 31954–31964 (2010).

53. Iwasaki, W. et al. Comprehensive Structural Analysis of Mutant Nucleosomes Containing Lysine to Glutamine (KQ) Substitutions in the H3 and H4 Histone-Fold Domains. Biochemistry 50, 7822–7832 (2011).

54. Hao, F. et al. Acetylation-modulated communication between the H3 N-terminal tail domain and the intrinsically disordered H1 C-terminal domain. Nucleic Acids Res. 48, 11510–11520 (2020).

55. Chittuluru, J. R. et al. Structure and nucleosome interaction of the yeast NuA4 and Piccolo– NuA4 histone acetyltransferase complexes. Nat. Struct. Mol. Biol. 18, 1196–1203 (2011).

56. Doyon, Y. & Côté, J. The highly conserved and multifunctional NuA4 HAT complex. Curr. Opin. Genet. Dev. 14, 147–154 (2004).

57. Balasubramanian, R., Pray-Grant, M. G., Selleck, W., Grant, P. A. & Tan, S. Role of the Ada2 and Ada3 Transcriptional Coactivators in Histone Acetylation. J. Biol. Chem. 277, 7989–7995 (2002).

58. Grant, P. A. et al. Yeast Gcn5 functions in two multisubunit complexes to acetylate nucleosomal histones: characterization of an Ada complex and the SAGA (Spt/Ada) complex. Genes Dev. 11, 1640–1650 (1997).

59. Kan, P.-Y., Lu, X., Hansen, J. C. & Hayes, J. J. The H3 Tail Domain Participates in Multiple Interactions during Folding and Self-Association of Nucleosome Arrays. Mol. Cell. Biol. 27, 2084–2091 (2007).

60. Gordon, F., Luger, K. & Hansen, J. C. The Core Histone N-terminal Tail Domains Function Independently and Additively during Salt-dependent Oligomerization of Nucleosomal Arrays. J. Biol. Chem. 280, 33701–33706 (2005).

61. Winkler, D. D., Muthurajan, U. M., Hieb, A. R. & Luger, K. Histone Chaperone FACT Coordinates Nucleosome Interaction through Multiple Synergistic Binding Events. J. Biol. Chem. 286, 41883–41892 (2011).

62. Wei, S., Falk, S. J., Black, B. E. & Lee, T.-H. A novel hybrid single molecule approach reveals spontaneous DNA motion in the nucleosome. Nucleic Acids Res. 43, e111–e111 (2015).

63. Tripathi, A., Tam, K. C. & McKinley, G. H. Rheology and Dynamics of Associative Polymers in Shear and Extension: Theory and Experiments. Macromolecules 39, 1981–1999 (2006).

64. Farr, S. E., Woods, E. J., Joseph, J. A., Garaizar, A. & Collepardo-Guevara, R. Nucleosome plasticity is a critical element of chromatin liquid–liquid phase separation and multivalent nucleosome interactions. Nat. Commun. 12, 2883 (2021).

65. Das, S. K. et al. Histone H3 Tail Modifications Alter Structure and Dynamics of the H1 C-Terminal Domain Within Nucleosomes. J. Mol. Biol. 435, 168242 (2023).

66. Shogren-Knaak, M. et al. Histone H4-K16 Acetylation Controls Chromatin Structure and Protein Interactions. Science 311, 844–847 (2006).

67. Di Pierro, M., Zhang, B., Aiden, E. L., Wolynes, P. G. & Onuchic, J. N. Transferable model for chromosome architecture. Proc. Natl. Acad. Sci. 113, 12168–12173 (2016).

68. Mirny, L. A., Imakaev, M. & Abdennur, N. Two major mechanisms of chromosome organization. Curr. Opin. Cell Biol. 58, 142–152 (2019).

69. Zhang, M. et al. Angle between DNA linker and nucleosome core particle regulates array compaction revealed by individual-particle cryo-electron tomography. Nat. Commun. 15, 4395 (2024).

70. Chahal, S. S., Matthews, H. R. & Bradbury, E. M. Acetylation of histone H4 and its role in chromatin structure and function. Nature 287, 76–79 (1980).

71. Antonin, W. & Neumann, H. Chromosome condensation and decondensation during mitosis. Curr. Opin. Cell Biol. 40, 15–22 (2016).

72. Iwasaki, W. et al. Contribution of histone N-terminal tails to the structure and stability of nucleosomes. FEBS Open Bio 3, 363–369 (2013).

73. Andresen, K., Jimenez-Useche, I., Howell, S. C., Yuan, C. & Qiu, X. Solution Scattering and FRET Studies on Nucleosomes Reveal DNA Unwrapping Effects of H3 and H4 Tail Removal. PLoS ONE 8, e78587 (2013).

74. Andrews, A. J., Downing, G., Brown, K., Park, Y.-J. & Luger, K. A Thermodynamic Model for Nap1-Histone Interactions. J. Biol. Chem. 283, 32412–32418 (2008).

75. Johnstone, C. P., Wang, N. B., Sevier, S. A. & Galloway, K. E. Understanding and Engineering Chromatin as a Dynamical System across Length and Timescales. Cell Syst. 11, 424–448 (2020).

