## Supplementary Figures for "Roles of histone chaperone Nap1 and histone acetylation in regulating phase-separation of chromatin arrays"

### Supplementary Information

Jia Gao[1], Hongyun Li[1], Song Tan[2], Ruobo Zhou[1,2], Tae-Hee Lee[1,*]

[1]Department of Chemistry and [2]Department of Biochemistry and Molecular Biology
The Pennsylvania State University, University Park, PA 16802

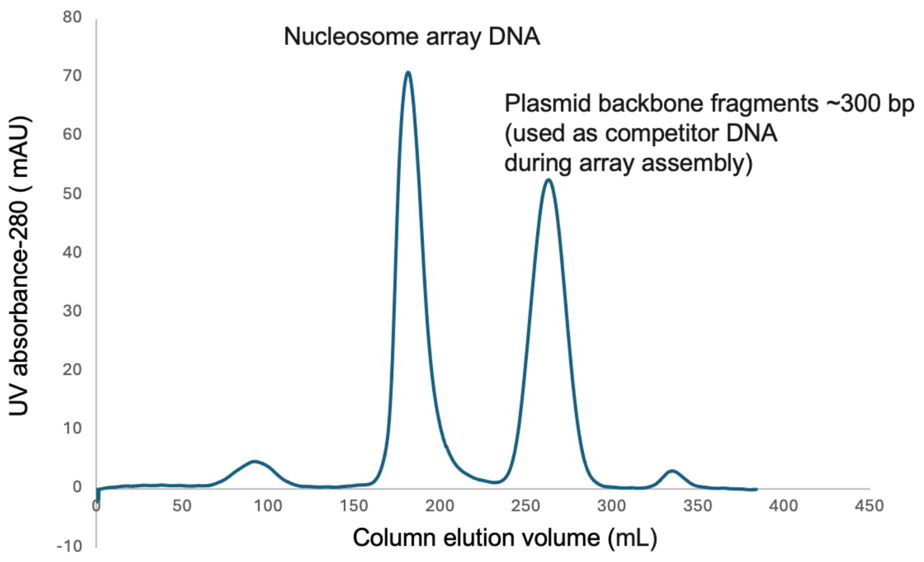


**Figure S1: A typical FPLC elution curve for DNA purification**

A Sephacryl S-500HR column (26 mm inner diameter, ~900 mm bed height, 0.7 mL/min elution rate) in conjunction with an AKTA Pure 25M (Cytiva) FPLC system was used to purify nucleosome array DNA after digesting plasmids. The first main peak represents the nucleosome array DNA. The second main peak represents the other digestion product which is ~300 bp in length. This shorter fragment is used as competitor DNA to “absorb” the leftover histones after saturating the nucleosome array DNA.

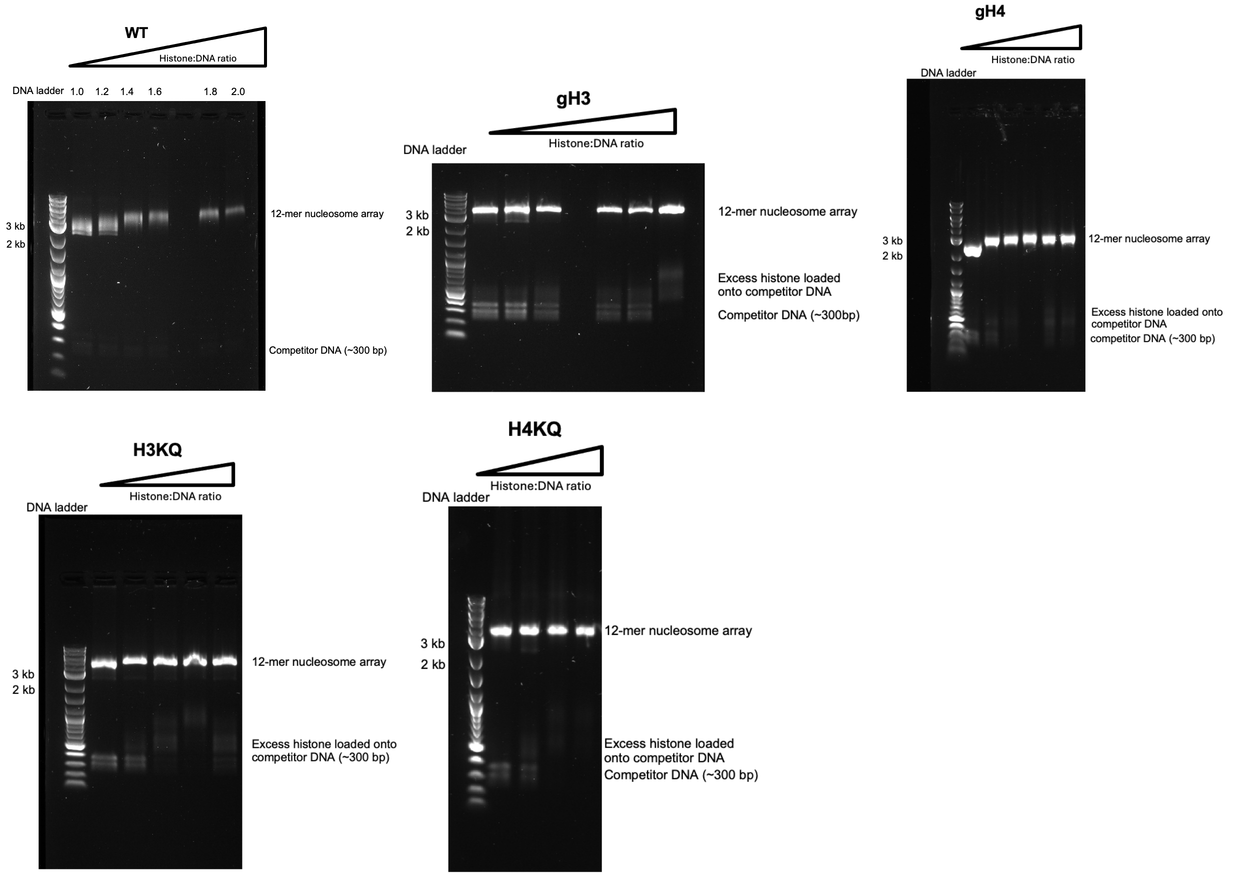


**Figure S2: Determination of DNA:histone ratio to saturate nucleosome array DNA for array preparation**

DNA:histone titration was carried out to determine the DNA:histone ratio to saturate the nucleosome array DNA with histones for the 5 types of samples (WT, gH3, gH4, H3KQ, and H4KQ). A DNA:histone ratio that shows both array saturation (i.e., no further upper shift in the gel) and a sign of histone loading on the competitor DNA without precipitation was selected and used for array preparation via dialysis, purification with FPLC, and analysis with AUC (Fig. S3). For the WT arrays, a transmission electron microscopic analysis has also been carried out to further confirm the assembly (Fig. S4).

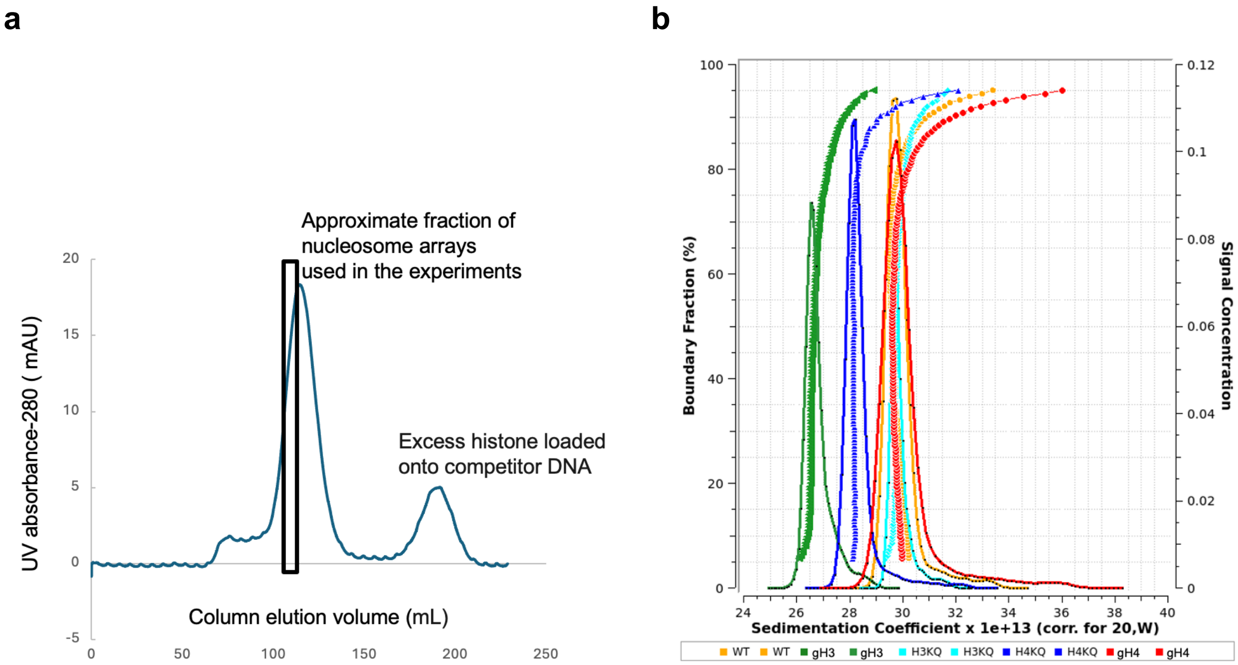


**Figure S3: A typical FPLC elution curve and sedimentation distributions of nucleosome arrays**

**a** A Sephacryl S-500HR column (Cytiva, 26 mm inner diameter, ~900 mm bed height, 0.3 mL/min elution rate) in conjunction with an AKTA Pure 25M FPLC system (Cytiva) was used to fractionate nucleosome arrays after saturating the DNA with histones in the presence of ~300 bp competitor DNA. The saturating ratio of DNA:histone was determined by native gel analyses as shown in Fig. S2. Fractions in the first peak were analyzed using analytical ultracentrifugation (AUC). The highest molecular weight fraction with minimal over-saturated arrays was used for the experiments. Its approximate elution volume is marked on the curve. **b** The AUC data for the FPLC fraction used in the experiments are shown for the 5 types of nucleosome arrays (WT, gH3, gH4, H3KQ, and H4KQ). The data were obtained with Optima Ultra Multiwavelength Analytical Ultracentrifuge (Beckman Coulter, analysis with absorption at 280 nm) and analyzed with UltraScan III (van Holde-Weischet plots for $S_{20,w}$ distributions).

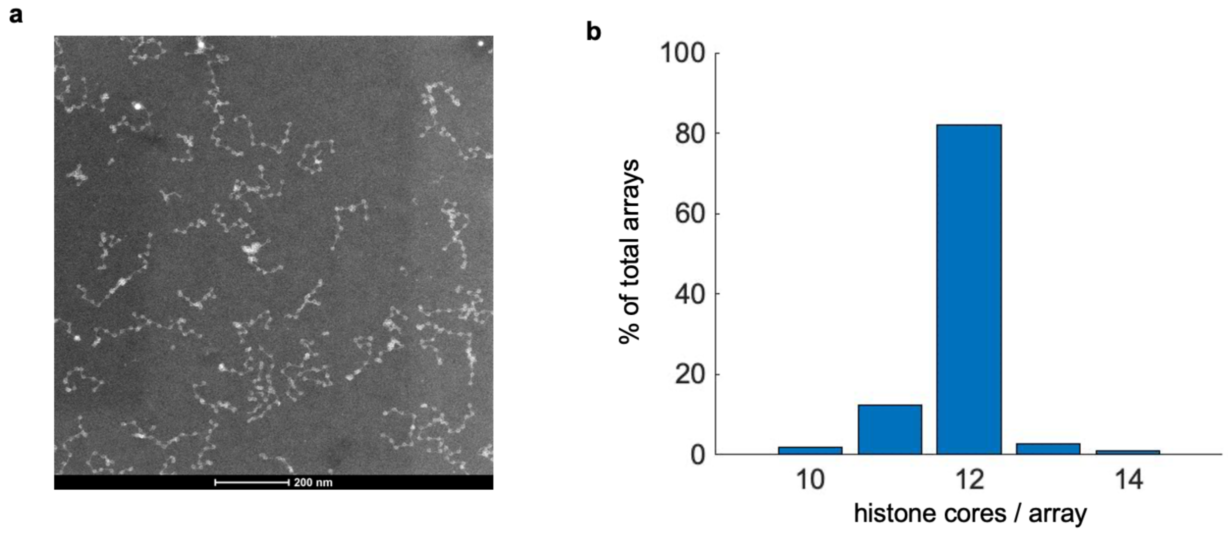


**Figure S4: A typical TEM image and a histogram confirming the proper assembly of nucleosome arrays**

**a** A transmission electron microscopy (TEM) image is shown to confirm nucleosome arrays made with unmodified histones (WT). **b** The number of histone cores was counted from all arrays that were visually resolvable. The histogram shows mostly 12-mer nucleosome arrays. The 11-mer arrays may have been over-counted due to the uncertainty associated with calling the number of histone cores based on visual inspection.

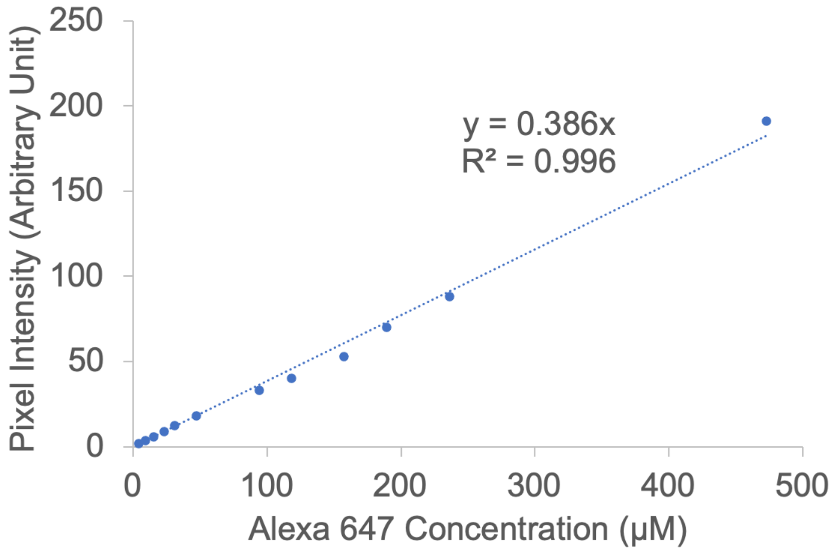


**Figure S5: Nucleosome concentration calibration**

Varying concentrations of the same fluorophore used to label histone (Alexa Fluor™ 647) were imaged with the same microscope (LSM880, Zeiss) used to carry out the FRAP measurements and the concentration measurements of nucleosomes within droplets. The pixel counts from a random region of interest at each Alexa Fluor 647 concentration were averaged to obtain the pixel intensity within the precision allowed by 3 significant figures. This calibration curve was used to convert the pixel intensity of a chromatin droplet to its nucleosome concentration. The nucleosome concentration was divided by 2 to give the final concentration shown in Fig. 4b as one nucleosome contains two Alexa Fluor 647 fluorophores labeled at two histone H4 molecules. The values are 326 ± 26, 521 ± 11, and 311 ± 33 μM for the WT, WT+Nap1, and H3KQ cases. The errors are measurement errors from multiple (n≥10) droplets.

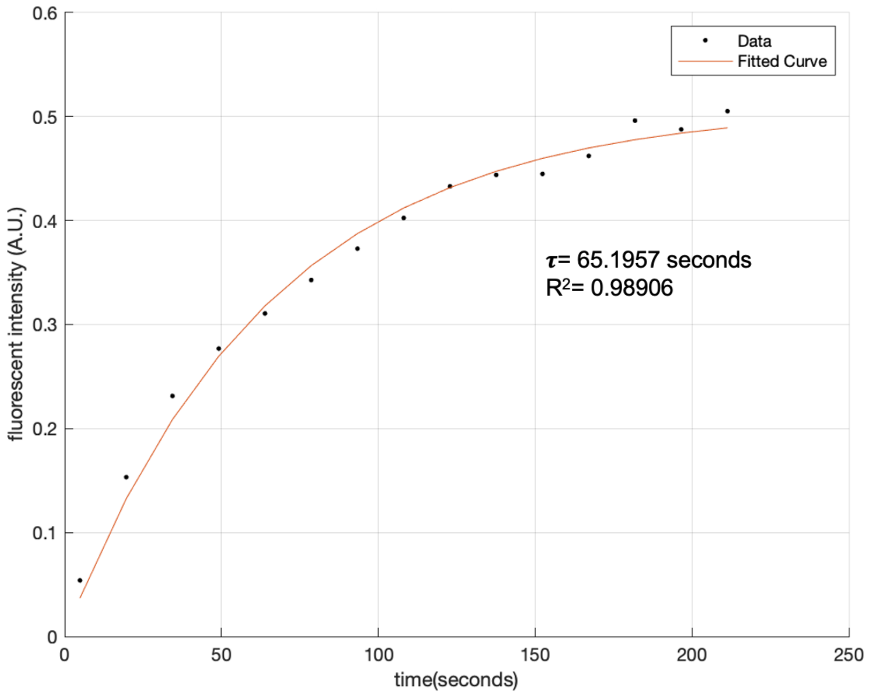


**Figure S6: Fluorescence recovery after photobleaching (FRAP) measurement**

A typical fluorescence recovery curve after photobleaching within a droplet (WT nucleosome arrays, labeled at H4 E64C with Alexa Fluor 647) shows fitting with an exponential recovery function described in Methods.
